# Analysis of Distributed Neural Synchrony through State-Space Coherence Analysis

**DOI:** 10.1101/2020.07.13.199034

**Authors:** Reza Saadati Fard, Kensuke Arai, Uri T. Eden, Emery N. Brown, Ali Yousefi

## Abstract

Established methods to track the dynamics of neural representations focus at the level of individual neurons for spiking data, and individual or pair of channels for local field potentials. However, our understanding of neural function and computation has moved toward an integrative view, based upon coordinated activity of multiple neural populations across brain areas. To draw network-level inferences of brain function, we propose a new modeling framework that combines the state-space model and cross-spectral matrix estimates – this is called state-space coherence (SSCoh). We define elements of the SSCoh and derive system identification and approximate filter solution for multivariate space processes. We expand SCoh for mixed observation processes, where the observation includes different modalities of neural data including local filed potential and spiking activity. Finally, we show an application of the framework to study neural synchrony across different brain nodes of a task participant performing Stroop task under different distraction levels.

## 1 Introduction

Numerous studies of brain activity, across a wide range of cognitive tasks and conditions, suggest evidence of precise and context-dependent temporal and modular organization of neural activity happening at multiple scales [1, 2, 3]. As a result, characterizing attributes of coordinated neural 1activity, or functional connectivity plays a critical role in advancing our understanding of healthy and pathological brain function [4, 5, 6]. Channel- and pair-wise analyses, which incorporate spectral power and coherence features, are widely used to characterize coordinated neural activity between brain areas nodes and associated brain functions [7, 8]. However, expanding this modeling framework to study population-wide associated brain activity is challenging, as evaluating statistical features of a distributed network is not straightforward using channel- or pair-wise measures. To build tools for network-level inference of the brain function, we require neural features representing population-wide associations of brain activity. Global coherence (GCoh), which is a more recent technique adopted for neural data analysis, provides a measure of overall synchrony across multiple brain areas [9, 10]. In GCoh, time-series data are transformed to the frequency domain using a Fourier transform and the cross-spectral matrix is estimated per each frequency. GCoh has been applied to neural data recorded during sleep-wake cycles, epilepsy, and anesthesia, and it has opened doors to better understand neural mechanisms distributed across brain areas under different cognitive states [10, 11]. However, during complex tasks and across changes in brain states, we expect different neural circuits to engage and disengage to facilitate multiple models of processing, leading to temporal dynamics in GCoh. To capture these changes, we propose a new modeling framework called state-space coherence (SSCoh). In SSCoh, the observation covariance – i.e., cross-spectral matrix, is defined as a function of the state process(es) to capture temporal covariation across multivariate time-series data. Such models are widely used in finance; however, the state-dependent cross-spectral analysis which is the core idea behind SSCoh is novel and provides a powerful tool for analysis of synchronous activity across high-dimensional time-series data. Here, we derive system identification and filter solution for the SSCoh. We expand the SSCoh for point-process and mixed observations, which address different modalities of neural data. We also discuss cross-frequency coherence analysis, which lets us study more complex network-level synchrony shaped by the interaction of networks at different frequencies. We show the application of SSCoh in a dataset where the neural activity of a subject is recorded by a 20-channel EEG device while performing a stress-induced Stroop task [12].

## 2 Methods

In this section, we define theories behind the state-space coherence and measures like GCoh which capture attributes of synchrony present in a multivariate time-series data. We start with the definition of GCoh and its estimation using the sample cross-spectral matrix [9]. We then introduce the covariate-dependent cross-spectral matrix solution to characterize synchrony in time-series data and develop a parameter estimation solution for this model. We expand the model to the broader SSCoh modeling framework, where the covariates in the cross-spectral matrix are latent and dynamical variables, we then derive the filter estimate and the parameter estimation solution. Finally, we discuss how the framework can be expanded to a broader class of neural data comprised of point-process observations or mixed neural data.

### 2.1 Coherence analysis in multivairate times series

The GCoh measure 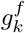 at frequency *f* and time interval *k* for a *M*-dimensional time-series data is defined by,

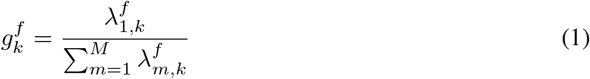

where, 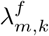 is the *m^th^* largest eigenvalue of the cross-spectral matrix calculated at time interval *k* [10]. The cross-spectral matrix, 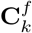, is defined by,

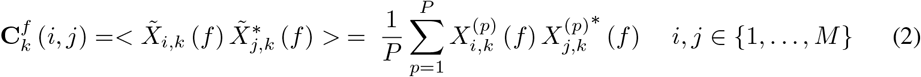

where 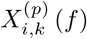 and 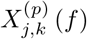 are *p^th^* estimate of Fourier transform (FT) of *i^th^* and *j^th^* channels at frequency *f* and time points [(*k −* 1) *N kN*]. *N* is the length of the window over which the FT is estimated and 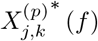 is the complex conjugate of 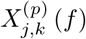 [10]. In practice, 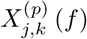 is a FT estimate of the tapered signal, where superscript *p* represents the *p^th^* of *P* different Slepian windows used to calculate tapered FT measurement. When the second-order stationarity assumption holds for multiple consecutive time intervals, the FT estimates from adjacents windows can be used in 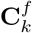 estimation [13].

The sample GCoh provides a measure of overall synchrony across multidimensional time series at a particular frequency. The 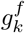 varies from 1/*M* to 1, where a larger value corresponds to more coherent activity across channels. We might be interested in how overall synchrony is reduced or increased during pathological versus healthy brain states, or during neuromodulatory or other environmental interventions. The GCoh measure allows these comparison. In the next section, we derive a regression model for the cross-spectral matrix to directly investigate the statistical relationship between these factors and GCoh by embedding the relationship between the time-series data and the possible underlying factors directly into the matrix definition.

### 2.2 Covariate-dependent coherence analysis

Under the covariate-dependent coherence model, we assume that the observation process, the FT measurements of different channels at frequency *f*, follow a multivariate complex normal distribution defined by,

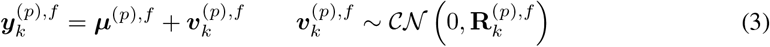

where, *k* represents the time interval, 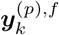 is the vector of *p^th^* tapered FT at frequency *f* of an *M*-dimensional signal. 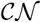 denotes complex normal distribution. Also, elements of ***μ***^(*p*),*f*^ and 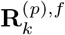 are complex variables. Under the assumptions of the central limit theorem, 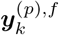 distribution follows a multivariate complex normal [14]; we assume the mean, ***μ***^(*p*),*f*^, is ergodic and the covariance matrix 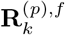 is defined as a function of covariates [15]. Note that under the mild wide-stationary assumption of time-series, the Fourier transform at different frequencies per each time interval become statistically independent; as a result, we build one model per each frequency. The covariance matrix 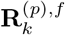 is a Hermitian symmetric matrix, which can be decomposed to,

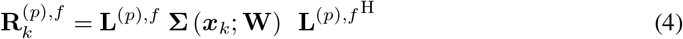

where **L**^(*p*),*f*^ is the eigenvector matrix and **Σ** is a diagonal matrix representing covariate-dependent eigenvalues [9]. ***x***_*k*_ is the *d*-dimensional vector of covariates at the time interval *k*, where each eigenvalue is defined as a function of the covariates and a set of free parameters, defined by weight matrix **W**, a *M* × (*d* + 1) matrix. The *m^th^*-eigenvalue is defined by,

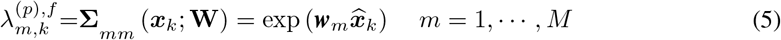

where 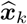 is [***x***_*k*_ 1] ^T^ and ***w***_*m*_ is the *m^th^*-row of **W**. We also assume the eigenvector matrix **L**^(*p*),*f*^ is a unitary matrix, an assumption which satisfies the following condition (note that this condition can be relaxed, which will be discussed later),

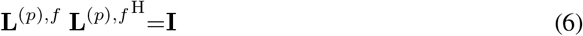

The GCoh for the model at time interval *k* is defined by,

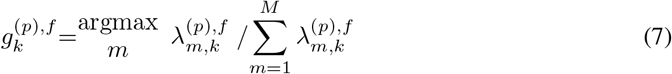

where 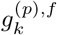 is the coherence measure for the *p^th^* taper at frequency *f*. The covariate-dependent coherence model free parameters include ***μ***^(*p*),*f*^, **W**, and **L**^(*p*),*f*^; these parameters define how the relationship between 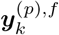 and ***x***_*k*_ get shaped per each time interval. To build any inferences given the covairates and time-series data, we must find maximum likelihood estimate of the model parameters. In the next section, we propose a solution to estimate the model parameters.

### 2.3 Parameter estimation

If the observation model parameters – **Θ** = {**W**, ***μ***^(*p*),*f*^, **L**^(*p*),*f*^ } – and covariates – ***x***_*k*_ – are known, the likelihood of observing 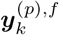 at time interval *k* is defined by,

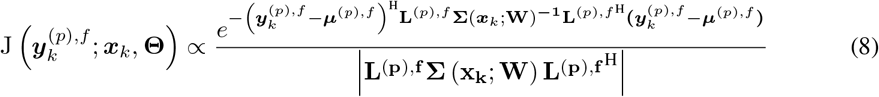

where J defines the likelihood at frequency *f* of 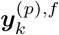 calculated using *p^th^* tapered FT measurement and |.| is determinant operator. Given **L**^(*p*),*f*^ is a unitary matrix, the likelihood function can be written as a function of the cross-spectral matrix eigenvalues,

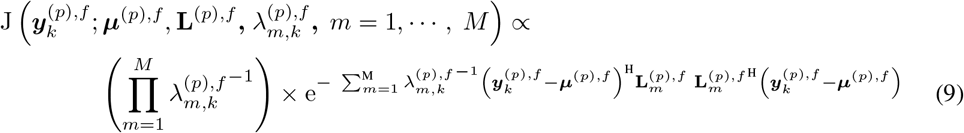

where 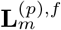 is the *m^th^* column of matrix **L**^(*p*),*f*^. Given Eq. 9, we suggest the Minorize-Maximzation algorithm [16] which provides a sequential update rule to find maximum likelihood estimate of the full likelihood function. We partition the parameter set **Θ** to {**W**} and { ***μ***^(*p*),*f*^, **L**^(*p*),*f*^ } sets. The parameter update for { ***μ***^(*p*),*f*^, **L**^(*p*),*f*^ } will be a constrained optimization defined by,

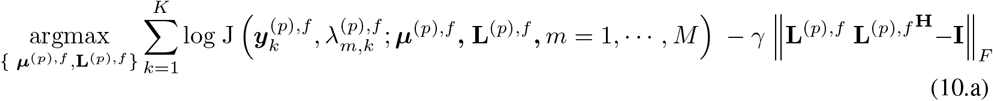

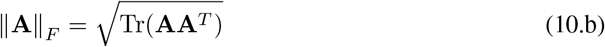

where ‖.‖_F_ in Eq. 10.b is Frobenius norm [17], and *γ* is the Lagrange multiplier. Given {**W**}, there is a closed form solution for { ***μ***^(*p*),*f*^, **L**^(*p*),*f*^ }.

To complete the update rule, we update {**W**} given the new updates for {***μ***^(*p*),*f*^, **L**^(*p*),*f*^}. The second optimization function is defined by

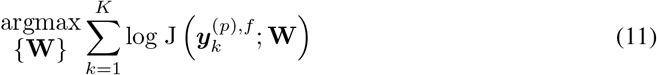

The full likelihood function is convex; as a result, we can find the optimal set of **Θ** with a recursive call of Eqs. 11 and 12.

Here, we built a parametric model of the cross-spectral matrix and demonstrated parameter estimation for the model. The ***μ***^(*p*),*f*^ corresponds to expected FT measure for frequency *f* over the whole time points, and **L**^(*p*),*f*^ defines eigenvectors – or eigenmodes – for synchronous activity across *M* channels. There is a broader class of problems, where the covariates are not accessible to a direct measurement, whilst the change of covariates over time will influence attributes of the synchrony across multivariate time-series data. For instance, the brain shows different levels of synchrony in particular frequency bands as a participant learning state evolves over time [18]. In the next section, we introduce SSCoh modeling framework, where the observation process is defined as a function of latent dynamical variables.

### 2.4 State-space coherence analysis

The state-space coherence model consists of two components, a state process and an observation process. We assume the state 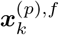, is a *d*-dimensional continuous latent variable where *f* represents a particular frequency, *k* represents the time interval index and *p* is the taper number. To characterize the state process dynamics over time, we have 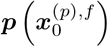 as an initial density function and a set of conditional density functions, 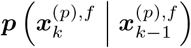, called the one-step transition densities. We assume a particular form for the one-step transition density, in which the distribution is Gaussian and the expected value of 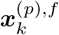 is a linear function of 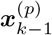 defined by,

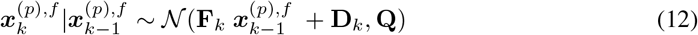

where 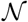 represents a *d*-dimensional multivariate Gaussian distribution, and the parameter set **Θ** consist of the state transition matrix **F**_k_, the drift input **D**_k_ and the *d* × *d* covariance matrix **Q**.

The observation process at time interval *k* is the tapered FT measure for all *M* channels. The observation process is defined by mean ***μ***^(*p*),*f*^ and state-dependent covariance noise 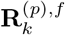:

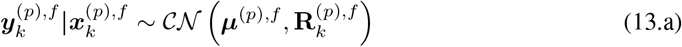

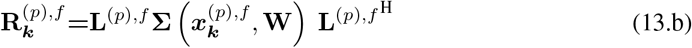

where 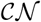 represents a complex *d*-dimensional multivariate Gaussian distribution, **L**^(*p*),*f*^ is the unitary eigenvector matrix, and **Σ** is the state-dependent covariance matrix.

The proposed model for the cross-spectral matrix assumes that eigenvectors are stationary over time and eigenvalues change as a function of the state process, assumptions reflecting new findings, where engagement and disengagement of different networks over time shape complex brain function [15]. Other forms of cross-spectral matrix decomposition like Bollerslev decomposition are used for time-varying covariance matrices [19], in which the correlation matrix is defined as a function of the state processes.

Figure 1 shows the block diagram of the SSCoh processing pipeline. In general, the state process might have more complex dynamics. For instance, they can have different forms of density functions or defined by both continuous and discrete processes. Discrete processes can address switching mechanisms like abrupt changes in the neural synchrony, evoked in response to a stimulus [20, 21, 22].

**Figure 1:**
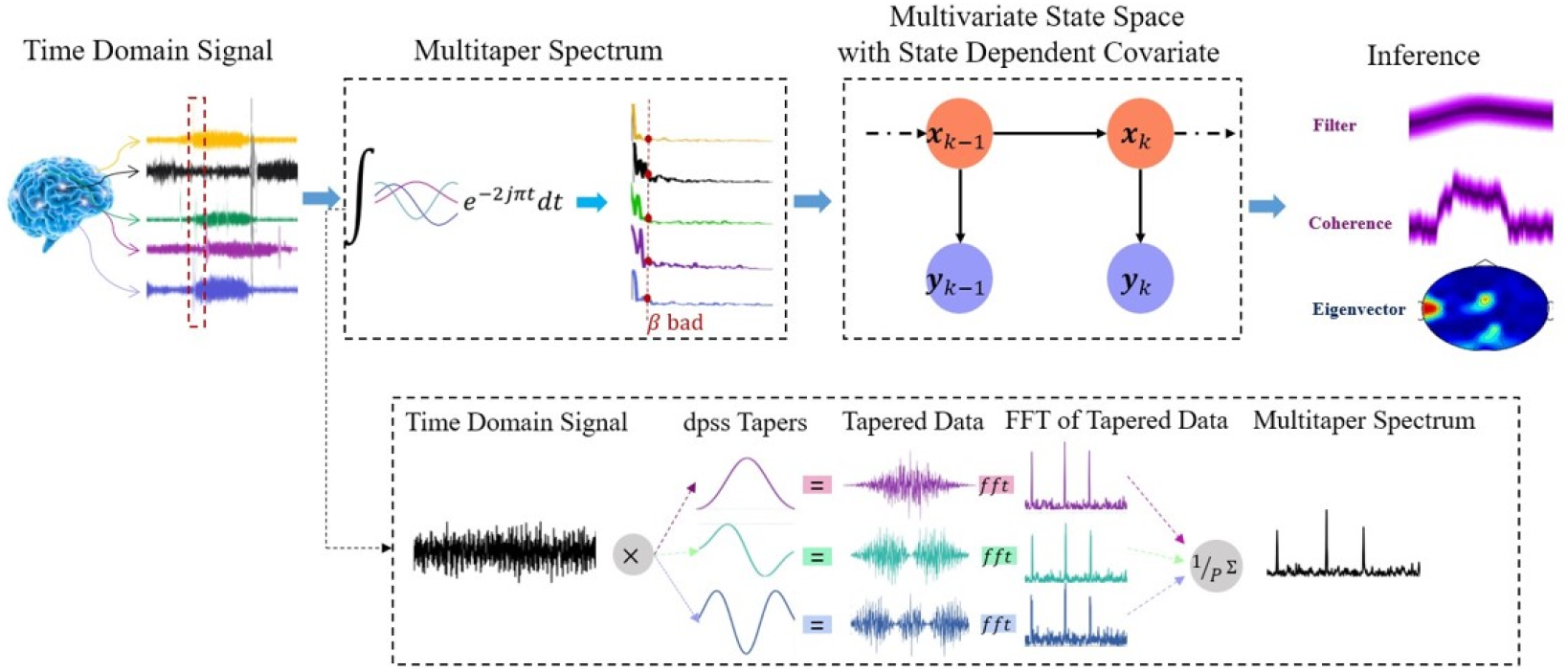
**SSCoh processing pipeline** time-series data of different locations of the brain captured by EEG electrodes. Multi-channel times series data is windowed and took tapered FT estimation. The observation has been made from the samples of the tapered spectral estimation. In the state-space model section, we run a state-space model in conjunction with the Expectation-Maximization algorithm to estimate parameters of the proposed model. Finally, in the Inference part, we would be able to obtain different measures of data including the distribution of coherency level

The core idea behind SSCoh is to move time-series data into the frequency domain to study the covariation. In finance, the idea of covariation in time across multivariate time-series are being embedded in the state-space models and has been used to address problems like stock volatility. For example, Multivariate Stochastic Volatility (MSV) model with a dynamic covariance model is used in optimal portfolio and risk management strategies [23, 24]. Similar solutions have been used to study associations across neural data in the time domain [15]. Despite the similarity between these models and SSCoh, SSCoh has a solid modeling foundation, provides a mechanistics understanding of network level neural synchrony, and the statistical properties of the observation are Gaussian and tractable. Also, analysis in the frequency domain is well aligned with our understanding of neural synchrony and gives a better picture of the brain functional dynamics.

Here, we described the SSCoh model. In the next sections, we will discuss the filter solution and parameter estimation for the SSCoh modeling framework.

### 2.5 Inference in SSCoh

We employ a Bayesian filter and smoother to find the posterior distribution of the state 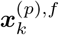 given the spectral measurements of the time-series data. Given the non-linear observation process, there is no closed-form solution for the filter and smoother. However, we can find the filter and smoother solutions using numerical, sampling, or approximate techniques [25, 26, 27]. Using the state posterior distribution, we can calculate the distribution of other metrics’ like GCoh or eigenvalues. Here, we propose a Gaussian approximation solution to derive a closed-form update rule for the posterior mean and covariance of the state, 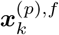. The update rules for mean and covariance are defined by

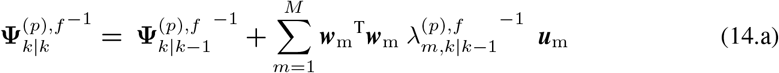

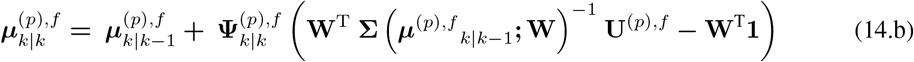

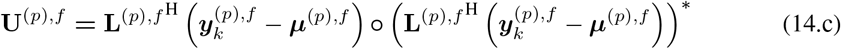

where 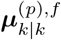 and 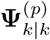 are the mean and covariance of posterior function and 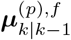 and 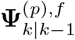 are the mean and covariance for one-step prediction for the *p^th^* Slepian taper FT at frequency *f*. 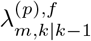 is the estimate of *m^th^* eigenvalue at time step *k* − 1, which is calculated from Eq. 5 with 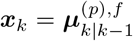. The ° represents the Hadamard product [28] and * is the conjugate operator. **1** is a vector of ones with length *M* and ***w***_*m*_ is the element of *m^th^* row of **W**. ***u***_*m*_ is the *m^th^* row of **U**^(*p*),*f*^ defined in Eq. 14.c In the Appendix A, we provide a detailed derivation of the solution that leads to Eq.s 14.a – 14.b.

The approximate solution can provide a reasonable approximation of the posterior distribution of the state processes; this is because, the posterior is unimodal and will have a shape close to multivariate normal. The approximation is fairly accurate, when the cross-spectral matrix has a sparse set of eigenvalues – here, we assume the time-series have finite energy.

Given the Eq.s 14.a – 14.c, we can use the Rauch-Tung-Striebel technique [29] to find the smoother estimate for the state processes. Using either filter and smoother, we can derive a closed-form solution for the expected eigenvalues and build other inferences like GCoh, phase delay across signal channels, or oscillation amplitude.

In the derivation of filter and smoother solutions, we assume the model parameters are known. In the next subsection, we discuss maximum likelihood estimation for the model parameters.

#### 2.5.1 Model parameter estimation

We use the Expectation-Maximization (EM) algorithm to find maximum likelihood estimates of the model parameters [30]. Under suitable conditions, the EM algorithm is guaranteed to converge to a local maximum of the full likelihood [31]. EM is run iteratively, where it increases the expected full log-likelihood per each iteration by updating the model parameters. The expectation is with regard to the posterior estimate of the state process given the whole observation; this posterior corresponds to the smoother solution described in the previous section. Each iteration of the EM algorithm consists of two steps: the expectation step (E-step), and the maximization step (M-step) [30]. In the E-step, the expectation of the full likelihood function is estimated given the posterior estimates of state calculated using current estimate of the model parameters. The E-step is defined by:

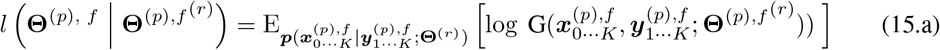

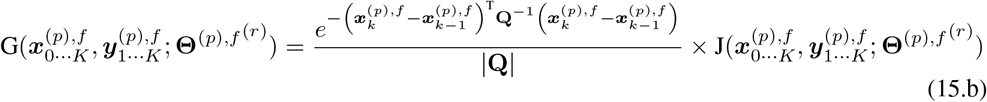

where E [.] is the expectation operator, and |.| is determinant operator, J is the likelihood function (defined in Eq. 8), G is the full-likelihood function, and 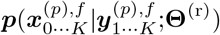 is the smoother distribution of 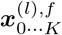 given the tapered FT measurement using current estimate of the model free parameters **Θ**^(r)^. *K* is the number of time intervals and superscript *r* represent the EM iteration.

In the M-step, the expected log-likelihood function is maximized by finding a new set of parameters. The maximization step is defined by,

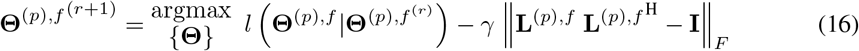

where the last term satisfies the model constraint which has been introduced in Eq. 6. **Θ**^(*p*),*f* (*r*+1)^ is the updated parameter set at iteration (*r* + 1). To find new updates for the model parameters, we require to calculate *l* (**Θ**^(*p*), *f*^ | **Θ**^(*p*),*f* (*r*)^) function first. This can be done use sampling techniques [32], where we draw samples from the states’ trajectory using the smoother estimation and becomes the maximization in Eq. 16 can be performed using numerical techniques like gradient ascent.

### 2.6 SSCoh for Point Process Data

The SSCoh model defined in Eq.s 12 and 13.a – 13.b assumes a continuous time-series data like local field potentials recorded from different brain areas as the input to FT. Non-continuous data modalities like population-level spiking activity can also be assessed for synchrony across different neural populations [33, 34]. We outline how SSCoh can be extended to point process observations [35, 36].

For point process data, several ways to estimate spike-train frequency content have been proposed [37, 38, 39]. Here, we use Mitra & Bokil [9]. Let us assume the *m^th^* channel of the multivariate time series data is a point process observation, its tapered FT measure at frequency *f* is defined by

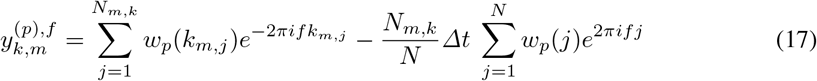

where *N*_*m*,*k*_ is the number of spikes for the [(*k* − 1) *N kN*] and *k*_*m*,*j*_ is time of *j^th^* spike in this interval for the *m^th^* channel. *w_p_* is the coefficient of *p^th^* taper and Δ*t* is sampling resolution.

Using Eq. 17, the SSCoh modeling solution can be applied to point process observations or mixed processes including both point processes and continuous observations.

## 3 Application

The data set that we have used is recorded by a 20-channel EEG device through performing a stress-induce Stroop task. In order to demonstrate the application of the proposed model, 5 channels have been chosen Figure 2 a). We hypothesize that two different state processes occur in different time scales and they can derive network level dynamics in the brain. The first state process represents the overall change in the neural dynamic and it would be able to capture slow changes in brain dynamics. The second one has been involved with shorter time scales and appeared in the model when we have in-congruent trial in the Stroop task. The relation between state processes and the observation can be expressed in the eigenvalues and Eq. 5 can be written as:

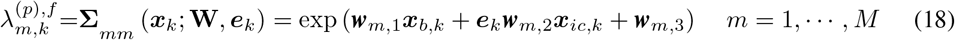

where ***x***_*k*_ is [***x***_*b*,*k*_ ***x***_*ic*,*k*_]^T^ in which ***x***_*b*_ and ***x***_*ic*_ represent the state process of baseline and incongruent task, ***w***_*m*,*n*_ is the element (*m*, *n*) of the matrix **W** and ***e*** is the indication function of in-congruent trial and has the value of 0 or 1; if we have an in-congruent trial at time interval *k* the value of ***e***_*k*_ would be equal to 1, else ***e***_*k*_ is 0.

**Figure 2:**
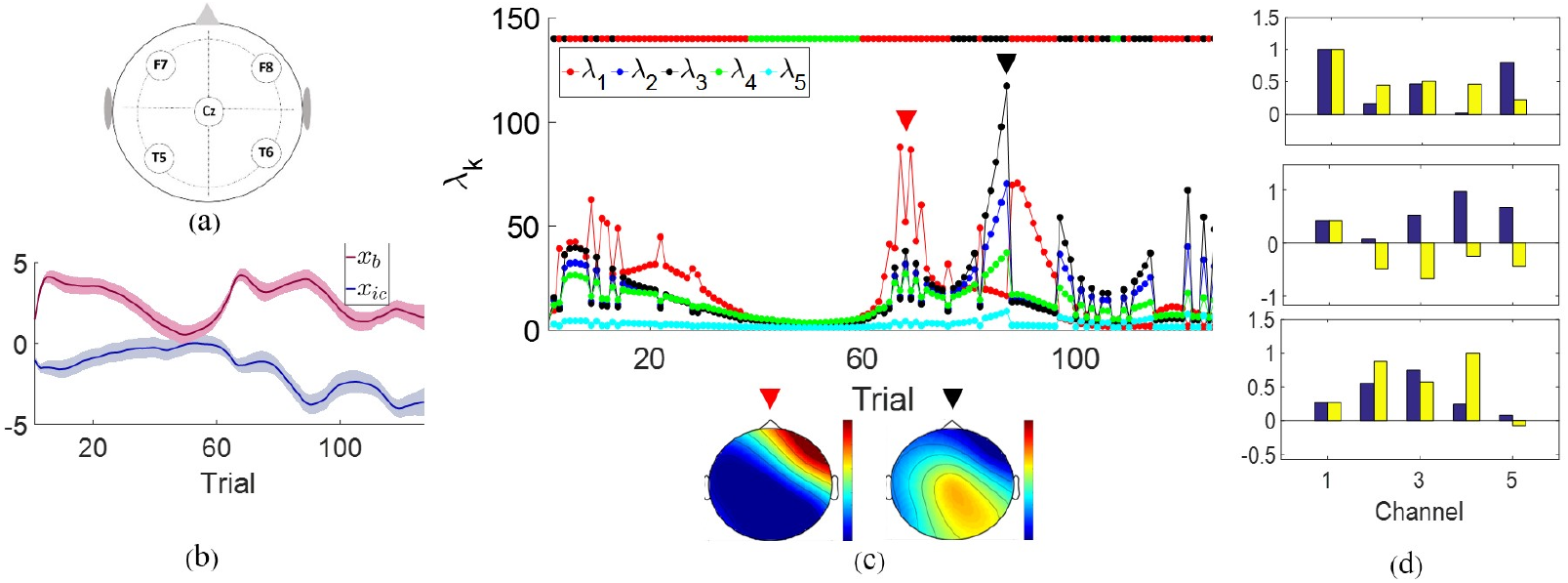
Modelling results in Stroop task. **a)** Positions of electrodes (F7, F8, Cz, T5, T6) being used in this analysis **b)** Smoother estimates for ***x***_*b*_ and ***x***_*ic*_ at the frequency of 6 Hz. The state estimate suggests clear network-level changes in the brain dynamics which is reflected in the domain eigenmode of neural activity. **c)** Mean of estimated eigenvalues per trial along with heatmap of the 2 dominant eigenvectors. There are 5 eigenvalues; the strength of eigenvalues changes as a function of the states. **d)** Initial and converged values of **W** matrix for the model. **W** is a matrix of 5 × 3.

Figure 2 shows the modeling results in a sample Stroop-task data set. Here, the model parameters including **W**, ***μ***, and state process noise are estimated using our proposed model identification solution – eigenvectors are fixed and estimated using sample cross-spectral analysis built over the whole experiment. The result suggests different eigenfunctions – specifically two dominant ones – emerge as the participants perform the task. The relative strength of the eigenvectors – which defines GCoh – changes throughout the experiment. For instance, from trials 30 to 60, GCoh is at its lowest level.

## 4 Conclusion

Here, we described theoretical foundations to capture network-level dynamics present through high-dimensional recordings. We defined the SSCoh framework and discussed how the relationship between the state processes and network-level neural activity can be characterized. We derived approximate filter and smoother solutions for SSCoh with a multi-variate state process and developed the parameter estimation algorithm. Through simulation results, we were able to validate filter and smoother results and also parameter estimation for W and ***μ***. Finally, we demonstrated the SSCoh application in a neural recording during a cognitive task. Through SSCoh, we can capture network-level brain dynamics at fine temporal resolution. Besides that, we can draw network-level inference which addresses the challenge of previous techniques developed based on pair-wise coherence. Despite its promising result, there are other questions which require to be addressed and resolved for SSCoh. For instance, we assumed a unitary eigenvector space, which might be a poor approximation in some cases, and a model parameter estimation procedure for the more general case needs to be explored. SSCoh can be expanded for cross-frequency analysis which is not discussed here. Stationary assumptions of the eigenvectors are not necessarily true, and we require incorporating changes in the eigenvector as a function of time or state process. Our future effort will focus on addressing these modeling questions and also combine SSCoh with behavioral readout to build a network-level neural encoder-decoder model.

